# Progressive behavioral and cognitive decline in *Drosophila* harboring *AD-*associated *APOE4* variants

**DOI:** 10.64898/2026.09.08.750261

**Authors:** Samaneh Biglari, Eshani Yeragi, David Bamisaye, Christian Gonzalez, Renata Garcia, Alexandra Losoya, Abhimanyu D. Arekere, Ciara Payne, Matthew J. Moulton, Alex C. Keene

## Abstract

Alzheimer’s disease (AD) is the most prevalent neurodegenerative disorder, and its incidence is rising rapidly with population aging. Pathologically, AD is characterized by the accumulation of amyloid-β (Aβ) plaques and hyperphosphorylated Tau neurofibrillary tangles. Human genomic studies have identified numerous risk alleles, with the APOE4 variant representing the strongest and most common genetic risk factor, present in approximately 75% of AD patients. However, APOE4 is neither necessary nor sufficient to cause disease, suggesting that additional genetic and environmental factors contribute to AD pathogenesis. Emerging evidence highlights a central role for oxidized lipid metabolism in AD. Disruption of lipid metabolism leads to lipid accumulation, reactive oxygen species (ROS) toxicity, and neurodegeneration, suggesting that oxidative stress may be a critical factor in enhancing AD susceptibility. To systematically investigate APOE function *in vivo*, we tested humanized *Drosophila* expressing the human APOE3, or APOE4 variants in place of the *Drosophila* ortholog *Glial Lazarillo* (*GLaz*). The lifespan of APOE3 and APOE4 flies do not differ under standard housing conditions, but the lifespan of APOE4 flies is significantly reduced when exposed to the ROS-promoting drug rotenone, supporting a multi-hit model of disease pathogenesis. APOE4 flies exposed to rotenone exhibit several AD-associated phenotypes, including age-related memory loss and chemosensory deficits, supporting the use of this model to investigate AD pathogenesis. Furthermore, progressive AD-associated phenotypes are also observed in APOE4 flies maintained on an obesogenic diet, suggesting that enhanced disease susceptibility is not specific to rotenone-induced stress but reflects a broader vulnerability to metabolic challenges. Together, these findings establish a scalable model to dissect APOE-dependent mechanisms and identify therapeutic targets in AD.

## Introduction

Alzheimer’s disease (AD) is the most common neurodegenerative disease, and its incidence is on the rise, especially in the US with a rapidly growing aging population ^1^. AD is pathologically differentiated from other neurodegenerative diseases by the accumulation of Aβ42 and hyperphosphorylated Tau into dense plaques (Aβ) and neurofibrillary tangles (Tau) in the brain ^2^. Genomic studies in humans have identified many allelic variants that associate with Alzheimer’s disease suggesting many unique genetic mechanisms contribute to the etiology of AD ^3^. The most common genetic variants associated with AD are in Apolipoprotein E (APOE), where humans carrying the APOE4 allele represent ∼75% of all patients ^4^. However, unlike many genetic variants associated with earlier-onset AD mutations, the APOE4 variant is neither necessary nor sufficient to induce AD^5^. The incomplete penetrance associated with APOE4 mutations has made systematically studying its effects in both animal models and humans more challenging. Therefore, developing *in vivo* models to investigate the cellular basis for APOE4-associated pathogenesis is critical for understanding the majority of AD cases.

APOE is a lipid-transport protein that is predominantly produced by astrocytes and facilitates the redistribution of cholesterol and other lipids between glia and neurons through interactions with LDL receptor family members ^6^. In addition to maintaining lipid and membrane homeostasis, APOE contributes to synaptic function, cellular responses to injury, and the clearance of cellular debris and amyloid-β (Aβ) ^6^. The APOE4 variant encodes a protein isoform that differs from the more common APOE3 isoform at amino acid 112 and confers substantially increased risk for AD. While the mechanisms linking APOE4 to AD pathogenesis remain incompletely understood, APOE4 is associated with altered lipidation and trafficking, impaired lipid homeostasis, and reduced Aβ clearance ^7^. These cellular changes are thought to contribute to increased Aβ deposition, neuroinflammation, endolysosomal and mitochondrial dysfunction, and synaptic deficits ^8^. Therefore, defining how APOE variants produce complex changes in cellular function and brain health is central to understanding AD pathogenesis.

The generation of *in vivo* mammalian models for the study of genetic variants, including in AD risk genes such as APOE and amyloid-β (Aβ), have advanced our understanding into the etiology and pathogenesis of neurodegenerative and other diseases ^9^. In many cases, this body of work has led to critical discoveries about the underlying biology of AD-based neurodegeneration, and a platform for testing pharmacological interventions ^10^. Nevertheless, the generation time and life expectancy of such models in the lab results in major bottlenecks in the ability of researchers to perform a systematic analysis of genetic risk factors or to screen compound libraries for therapeutic treatments in a feasible timeframe ^11,12^. It has been particularly challenging to assess the combinatorial effects of multiple genetic risk factors and the effects of environment on genetic risk.

Studies in animals and human tissue demonstrate the consequences of accumulated oxidized lipids in the pathogenesis of AD ^13,14^. In response to cellular stress, neurons produce and then secrete oxidized lipids via ABCA transporters. These lipids are then loaded onto the apolipoprotein, APOE, to form lipoprotein complexes ^15^. Glial cells then endocytose these lipidated APOE molecules and upon degradation of the complexes in lysosomes, the lipids are transferred to the ER where lipid droplets form, thus sequestering these toxic, oxidized lipids. In this manner, glia play a critical role in preventing ROS-induced neurotoxicity since loss of proteins required for lipid droplet formation, including APOE and ABCA1 and 7 orthologs as well as several other key proteins required for endocytosis, results in a failure to accumulate lipid droplets into glia leading to neuronal dysfunction and death ^7^. Therefore, understanding the role of glia in sequestering oxidized lipids is critical for understanding the role of APOE4 in AD pathogenesis.

The fruit fly, *Drosophila* melanogaster, provides a powerful model to study Alzheimer’s disease ^16,17^. *Drosophila* exhibit complex behaviors including forming associative memories and sleep, both of which are mediated through conserved genetic and neural mechanisms ^18,19^. Expression of AD-associated Aβ variants, including the highly pathogenic Arctic variant, results in neurodegeneration accompanied by deficits in sleep, memory, sensory processing and reduced longevity, phenocopying the human disease ^20–24^. In addition, these models have been used to test the effects of numerous pharmacological agents on the disease, as well as the effects of AD variants on defined cell-types ^25,26^.

The genetic amenability of *Drosophila* provides the ability to test the effects of specific human variants on AD-associated phenotypes. We have developed humanized flies expressing distinct variants of APOE that include common APOE3, or AD-associated APOE4 in substitution of the *Drosophila* APOE functional analog, *GLaz* ^27,28^. These models take advantage of the T2A-GAL4 system, in which the coding sequence of the yeast transcription factor GAL4 is inserted into the *GLaz* locus ^29^. This insertion disrupts endogenous GLaz function while driving GAL4 expression in the same spatiotemporal pattern as GLaz. These flies are crossed to UAS-APOE variant lines (GLaz^T2A-GAL^^4^ > UAS-hAPOE), generating animals that express human APOE under the control of the endogenous GLaz regulatory elements. Previous characterization of these flies suggests the neurotoxicity associated with APOE4 variants results in lipid accumulation that contributes to pathogenesis, consistent with the well-established accumulation of lipids in AD ^27^. However, the broader effects of the APOE4 variant on longevity and other AD-associated phenotypes have not been investigated. The development of this model that phenotypes the human disorder provides the opportunity to systematically investigate the effects of APOE on disease pathogenesis.

Here, we systematically investigated the effects of human APOE variants on Drosophila physiology and pathogenesis. We found that the consequences of APOE4 expression are exacerbated by exposure to the reactive oxygen species (ROS)-inducing toxin rotenone, supporting a multi-hit model of neurodegenerative disease. APOE4-expressing flies exposed to rotenone exhibited deficits in learning and chemosensory processing, along with reduced lifespan. Furthermore, APOE4 flies displayed increased sensitivity to both environmental and metabolic challenges, including rotenone exposure and an obesogenic diet, suggesting reduced physiological resilience to diverse stressors. Collectively, these findings demonstrate that humanized APOE4 flies recapitulate multiple phenotypes associated with Alzheimer’s disease and provide a tractable model for investigating the molecular mechanisms and behavioral consequences underlying disease pathogenesis.

## Results

To investigate the effects of APOE variants on longevity we compared longevity in humanized APOE3 and APOE4 flies where the APOE variant is selectively expressed in place of its *Drosophila* ortholog *GLaz* ^27^. We compared flies of both genotypes housed on standard food to those housed on standard food containing a 50μM concentration of the ROS-inducing drug rotenone ^30^. Flies were placed in the Drosophila Activity Monitoring System (DAMS) one day following eclosion. Locomotor activity was continuously monitored by recording infrared beam breaks until the time of death (Fig 1A) ^31^. There was no significant difference in longevity between APOE3 and APOE4 flies housed on standard food (Fig 1B). While rotenone reduced longevity in both APOE3 and APOE4 flies, lifespan was significantly shortened in APOE4 flies fed rotenone compared to vehicle-fed flies, suggesting enhanced sensitivity to rotenone in flies harboring humanized APOE4 (Fig 1B). To validate these findings under social housing conditions, we measured longevity in flies maintained in groups of 10 males per vial and monitored survival. Similar to flies tested in isolation, longevity of APOE4 flies fed rotenone was significantly less than all other groups tested (Fig 1C). Under social conditions, APOE4 flies lived significantly shorter than APOE3 counterparts, raising the possibility that social experience and housing condition impact the effect of APOE alleles and rotenone exposure. Together, these findings suggest APOE4 robustly affects resilience to ROS stress, providing a multi-hit model of AD in humanized *Drosophila*.

**Figure 1.**
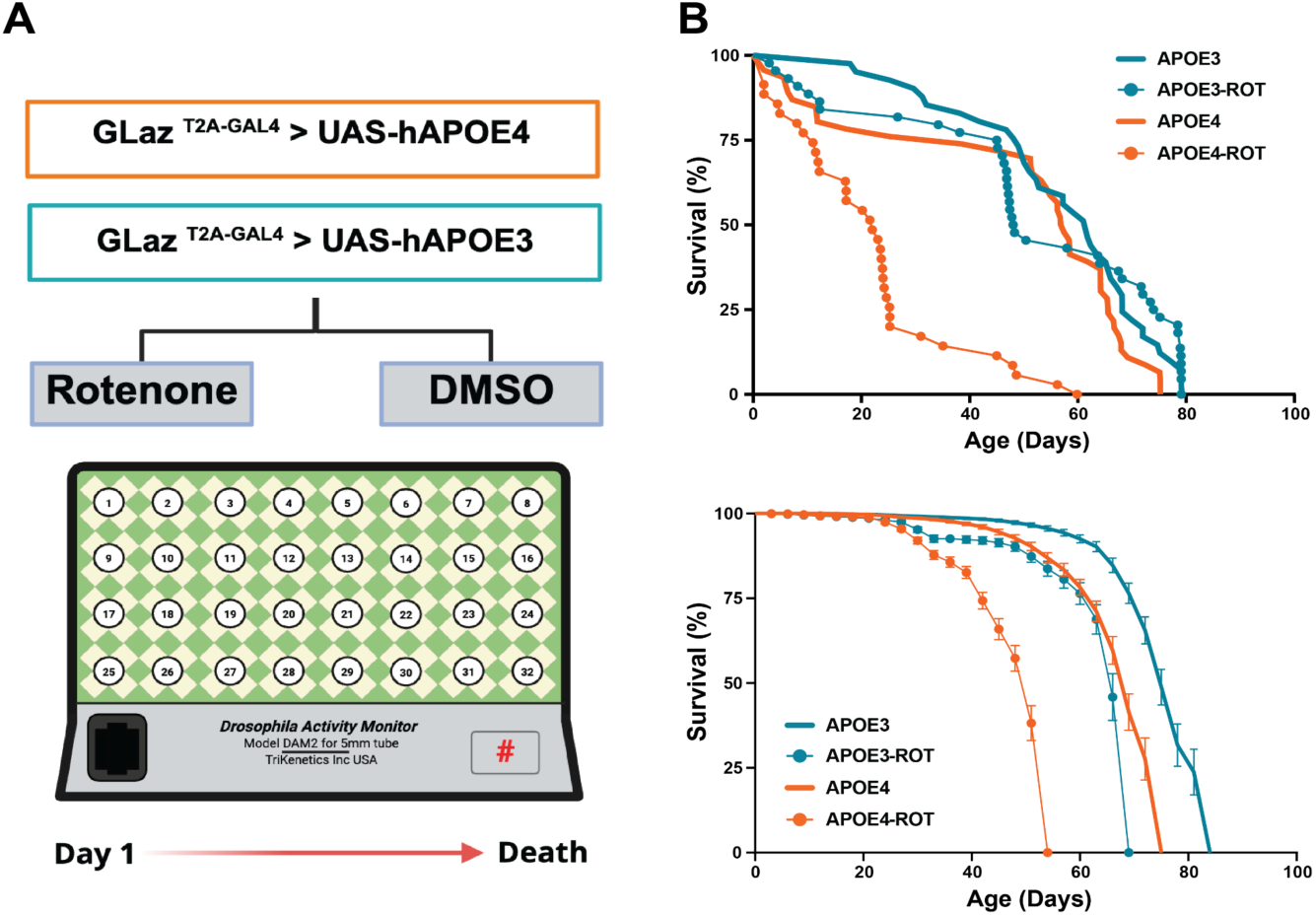
Rotenone exposure alters survival in flies expressing humanized APOE3 or APOE4. **(A)** Schematic overview of the experimental paradigm. One day old male flies expressing APOE3 or APOE4 were maintained on food containing rotenone or vehicle control (DMSO) and survival was assessed across groups. **(B)** Survival of individually housed flies tracked across lifespan using Drosophila Activity Monitor (DAM) system revealed significant reduction in longevity under rotenone containing food in APOE3 and APOE4 flies, with the shortest lifespan observed in APOE4 rotenone fed flies (log-rank Mantel-Cox test, χ²(3) = 81.28, *p* < 0.001). **(C)** Survival of flies socially housed differed significantly across genotypes with reduced longevity on APOE4 flies fed rotenone containing food (log-rank Mantel–Cox test, χ²(3) = 534.7, *p* < 0.001).

Sleep is disrupted in Alzheimer’s patients, and a number of reports suggest sleep is disrupted in AD model flies ^20,23,32,33^. To determine whether sleep is disrupted in APOE4 flies we quantified sleep across the lifespan. When comparing sleep across the lifespan, we found that sleep duration was relatively stable across the lifespan (Fig S1A). However, when focusing specifically on young flies, when sleep has previously been studied in AD model flies, we found that sleep was significantly lower in APOE4 flies, compared to APOE3 flies (Fig S1B). Treatment with rotenone did not affect sleep duration in APOE3 flies but significantly increased sleep duration in APOE4 flies, potentially reflecting increased sleep need or reduced sleep quality (Fig. S1B). To determine whether these effects reflected changes in sleep specifically or a more general alteration in locomotor activity, we measured waking activity. Rotenone had no effect on waking activity in APOE3 flies but induced hyperactivity in APOE4 flies (Fig S1C). Together, these findings support previous findings indicating sleep differences in AD-model flies.

It is possible that APOE3 and APOE4 flies consume different amounts of food, confounding interpretations about the effects of rotenone. We used the blue dye assay to compare feeding in APOE3 and APOE4 flies. Briefly, starved flies were housed on food containing blue dye for 30 minutes ^34,35^ (Fig S2A) and then were flash frozen and spectrophotometry was used to quantify the amount of food. At 5 days, no significant differences were detected between any of the groups tested (Fig S2B). At 20 days, there was a lower food consumption in rotenone fed APOE4 flies compared to APOE3 flies on rotenone, though no significant differences were detected between DMSO and rotenone treated flies within either APOE3 or APOE4 genotypes (Fig S2C). Therefore, the effects of rotenone on longevity and other traits are unlikely to be attributed to changes in feeding or increased rotenone intake in APOE4 flies.

Reduced chemosensory processing is a hallmark of many neurodegenerative diseases including Alzheimer’s ^36^. *Drosophila* exhibit a robust Proboscis Extension Reflex (PER) in response to an appetitive tastant, providing a quantifiable measure of feeding propensity ^37^. We have previously shown that PER is diminished with age, and in flies expressing pathogenic Aβ42 ^24^. To examine the relationship between aging and taste in APOE flies, we measured PER in response to varying concentrations of fructose in young (5 day) and aged (20 day) flies. In this assay, applying a tastant directly to the proboscis evokes a reflexive response that does not depend on post-ingestive feedback ^38,39^ (Fig 2A). To define the sensitivity to gustatory stimuli, flies were provided with different concentrations of fructose ranging from 1 mM to 1 M. In 5-day-old flies, the response to these concentrations ranged from 15% to 100%, providing a dose-response readout of sensory-response threshold. At 5 days, rotenone reduced PER overall. However, there was no effect of rotenone treatment on APOE genotype (Fig 2B). Unlike the younger flies, at 20 days rotenone’s effect depends strongly on APOE genotype, with a near complete loss of taste response in flies fed less than 50 mM fructose. (Fig 2C). Quantification of the response to 50mM fructose reveals no loss of PER in 20-day old APOE3 flies, and a significant reduction in PER in 20-day old APOE4 flies fed rotenone (Fig 2D). Therefore, the APOE4 allele is associated with reduced gustatory sensitivity, and reduced resilience to rotenone treatment.

**Figure 2.**
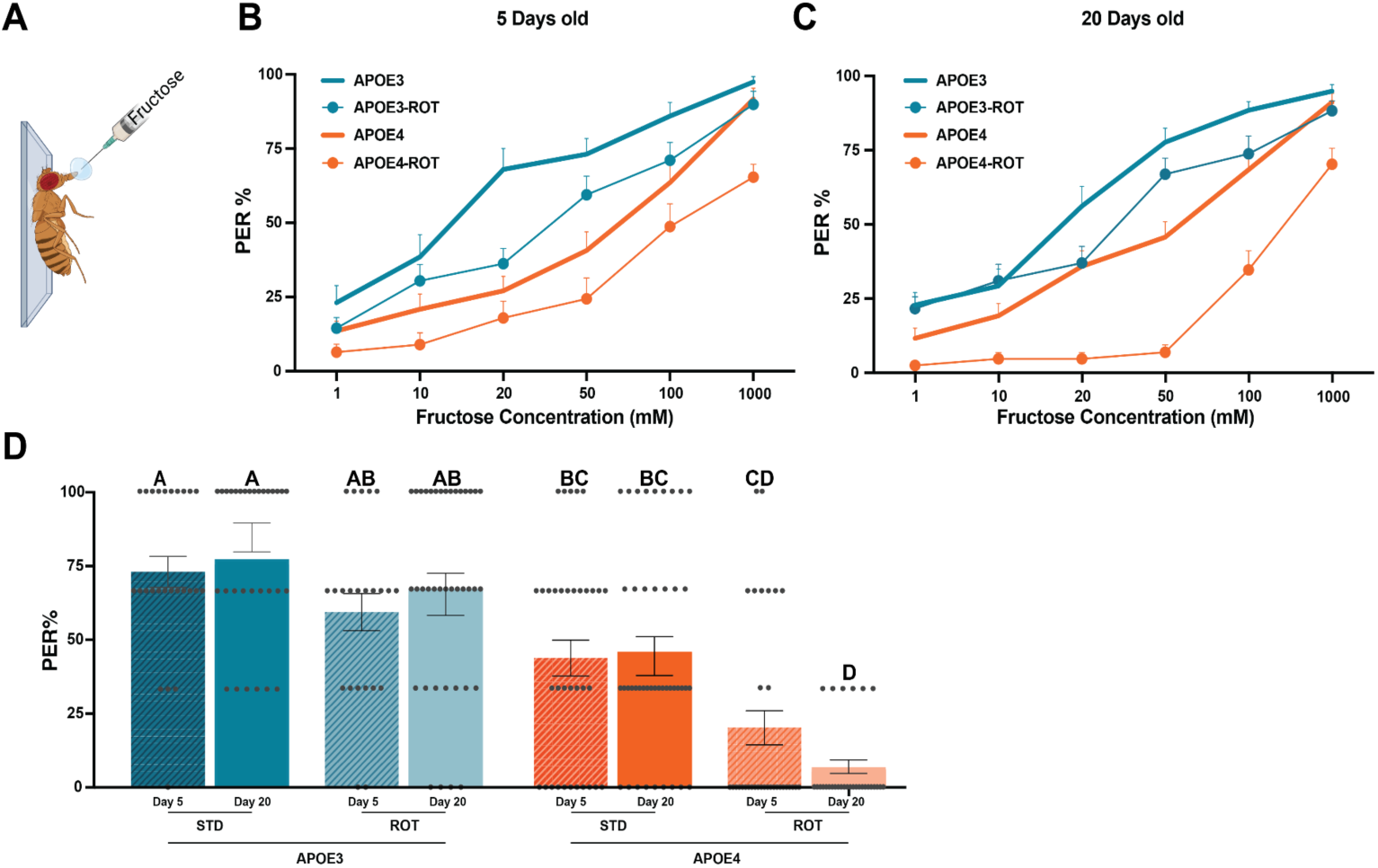
APOE4 enhances rotenone-induced impairment of fructose responsiveness in 20-day-old flies. **(A)** Proboscis Extension Reflex (PER) was measured in flies after 14 h of starvation. **(B)** In 5 day old flies, there were no significant differences between the groups (*p* = 0.934) (*n* > 26) **(C)** In 20 day old flies, there was a significant effect between APOE and rotenone (*p* < 0.001). Šídák’s multiple comparisons test showed that rotenone did not significantly alter PER in APOE3 flies (p = 0.884), while strongly reduced PER in APOE4 flies (*p* < 0.001) (*n* > 30). **(D)** Comparison of responses at 50 mM fructose shows that PER in 20-day-old rotenone fed APOE4 flies was significantly lower than APOE3 and standard fed APOE4 flies (*p* < 0.05). Groups not sharing a letter differ at adjusted *p* < 0.05. Individual points represent flies and bars represent mean ± SEM.

In addition to taste, flies display robust olfactory behaviors that diminish with age ^40^. We sought to determine whether the deficits in APOE4 chemosensation generalize to olfactory behavior ^41^. We tested flies in the olfactory-trap assay that measures their ability to select attractive odor sources ^42^. Briefly, flies were placed in a bottle and allowed to enter one of two traps through a one-way funnel: a control trap containing DI water or an experimental trap containing the attractive odorant acetic acid (Fig S3A). There was no significant decrease in olfactory performance of APOE3 flies, while preferential selection of acetic acid was abolished at 20 days old in APOE4 flies fed standard food or rotenone (Fig S3B). Therefore, age-related loss of performance is identifiable in APOE4 flies, and exacerbated by rotenone feeding.

*Drosophila* form robust associative memory to a variety of stimuli ^43,44^. Across many different memory paradigms, memory diminishes with age, and in neurodegeneration models ^45–49^. To determine whether age-related memory decline is exacerbated in APOE4 model flies, we tested these groups in an aversive taste memory assay ^50^. Briefly, pairing sucrose presentation to the tarsi with bitter quinine presentation to the proboscis results in the formation of aversive taste memories where flies suppress response to the presentation of sucrose alone ^51,52^ (Fig 3A). At five days of age, no differences were detected between any of the groups indicating that memory is intact at a young age, and that flies are not acutely sensitive to rotenone exposure (Fig 3B). Conversely, at 20 days old, taste memory was reduced across the last two training and test trials in rotenone fed APOE4 mutant flies compared to APOE4 flies housed on normal food and both groups of APOE3 flies, while there were no differences in training between flies (Fig 3C). At 20 days old memory was not impacted by rotenone in APOE3 flies but was significantly reduced in APOE4 flies housed on rotenone compared to APOE4 flies housed on normal food and both groups of APOE3 flies (Fig 3D). Therefore, APOE4 accelerates memory loss, and this increases with ROS exposure.

**Figure 3.**
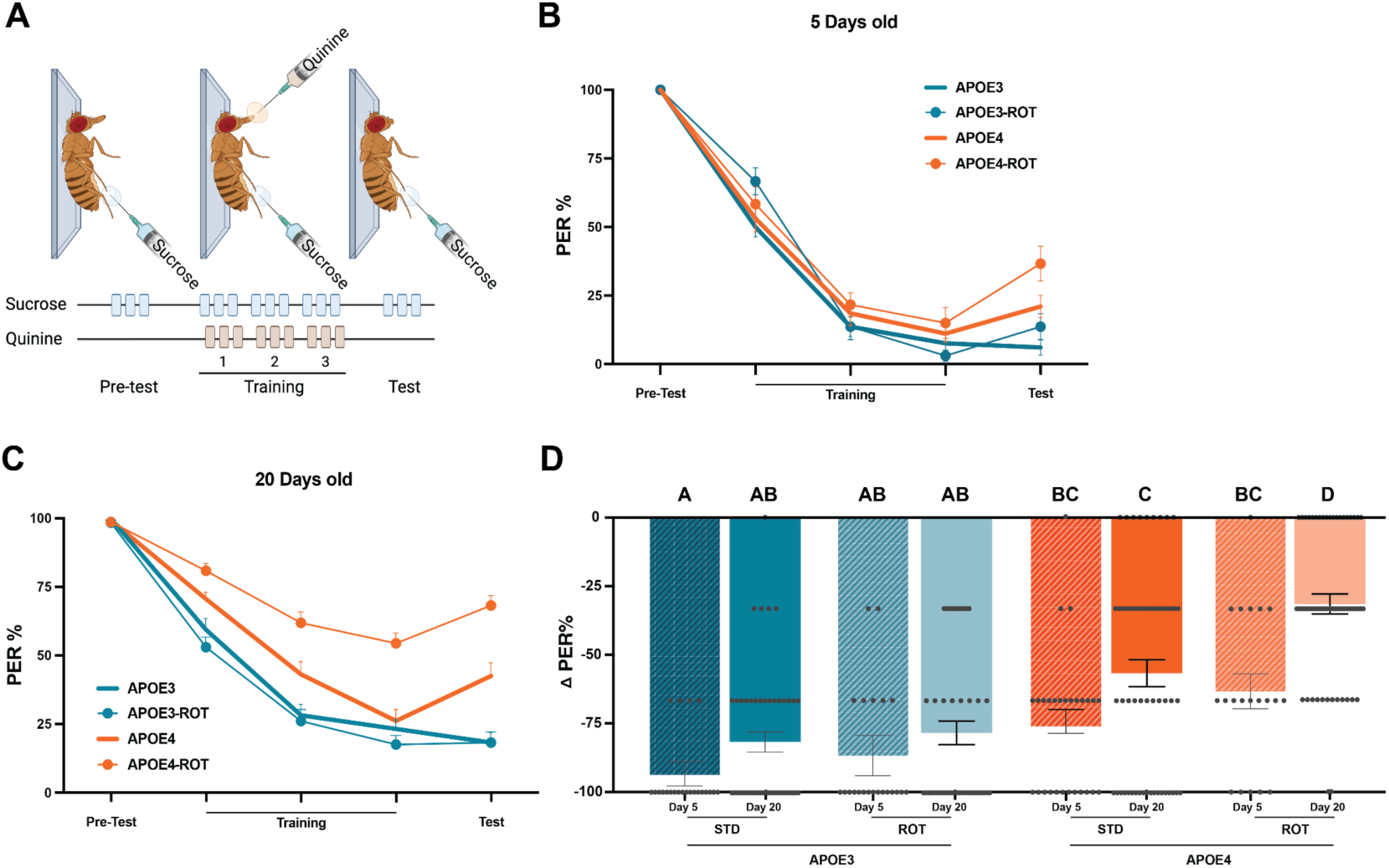
Aging and rotenone exacerbate deficits in aversive learning and memory in APOE4 flies. **(A)** Schematic of aversive taste memory assay. PER was measured in male flies. **(B)** In 5-day-old flies no differences were detected between the groups. (*n* > 20). **(C)** At 20 days of age, rotenone fed APOE4 mutant flies showed impaired taste memory during the final two training trials and subsequent test trials compared with all other groups (*p* < 0.05) (*n* > 47). **(D)** Quantification of aversive taste memory performance revealed that at 20 days of age, rotenone treatment did not affect memory in APOE3 flies, whereas memory was significantly impaired in rotenone fed APOE4 flies compared with APOE4 flies on standard food (*p* = 0.0008) and both APOE3 groups (*p* < 0.0001). Groups not sharing a letter differ at adjusted *p* < 0.05. Data are shown as mean ± SEM.

Aging in *Drosophila* is associated with a decline in intestinal stem cell homeostasis, leading to disruption of gut barrier integrity, which has been proposed to contribute to age-related mortality ^53,54^. In *Drosophila*, intestinal barrier integrity can be assessed by feeding flies blue dye, and visualizing whether it localizes to the gut, indicating an intact gut, or is present throughout the body cavity, revealing gut permeability (Fig 4A) ^55^. We compared gut permeability in young and old APOE3 and APOE4 flies with and without rotenone (Fig 4B). Consistent with previous reports, there were no examples of gut permeability in young flies across any of the groups tested (Fig 4C). Further, there was no detectable gut permeability in APOE3 flies fed rotenone at either age, however, there was significant gut permeability in 20-day old APOE4 flies fed rotenone. Therefore, rotenone increases gut permeability in aged APOE4, but not APOE3 flies.

**Figure 4.**
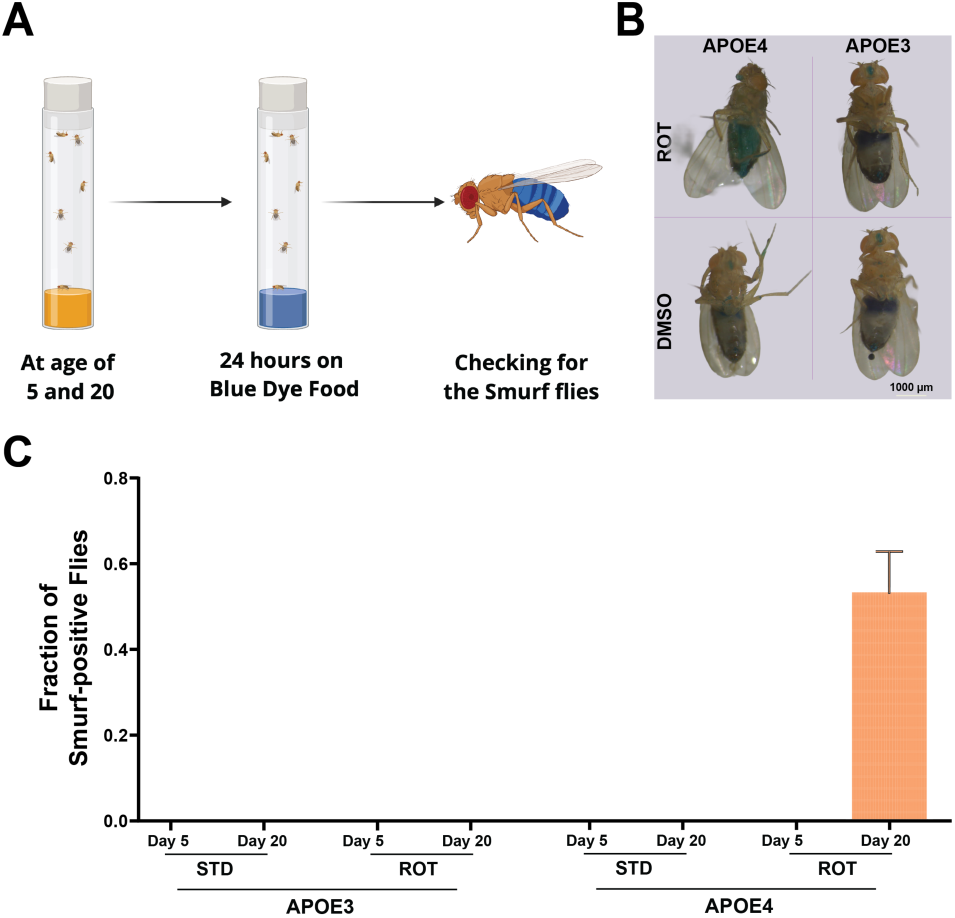
Rotenone induces age-dependent intestinal barrier permeability in APOE flies. **(A)** Schematic of the gut-permeability assay. Smurf assay used to assess intestinal barrier dysfunction in 5 and 20-day old APOE3 and APOE4 flies maintained on food containing rotenone or vehicle control (DMSO). **(B)** Representative images for each group tested at 20 days of age. Non-Smurf flies retained the blue dye within the gastrointestinal tract, whereas Smurf-positive flies showed blue dye leakage into the surrounding body cavity, indicating loss of intestinal barrier integrity. **(C)** Aged APOE4 flies (20-day-old) fed rotenone had a significantly greater fraction of Smurf-positive flies (*p* < 0.001), while the remaining groups showed no Smurf-positive phenotype (*n* = 35).

Many environmental and lifestyle factors influence AD risk. Notably, obesity and type 2 diabetes are among the strongest modifiable risk factors, highlighting a potential role for metabolic dysfunction in disease pathogenesis ^56^. Feeding *Drosophila* a high sugar diet (HSD) induces obesity and insulin-resistance phenocopying many aspects of diabetes^57,58^. To examine the effects of HSD on APOE-model flies we compared longevity of APOE3, and APOE4 flies reared on standard diet or HSD (Fig 5A). In agreement with previous data, there was no difference in longevity between APOE3, and APOE4 flies housed on a standard diet (Fig 5B). Lifespan was shortened in both APOE3 and APOE4 flies housed on HSD yet was significantly shorter in APOE4 flies than APOE3 flies, suggesting increased sensitivity to HSD (Fig 5B). Sleep was significantly increased on HSD in APOE3 and APOE4 flies as compared to the standard food (Fig S4A-B). To determine the effect of HSD on chemosensation, we examined responsiveness to fructose (Fig 5C). At 5 days of age there was a significant difference between APOE4 flies fed HSD compared to all other groups (Fig 5D). At 20 days of age, there were no differences in APOE3 and APOE4 flies housed on standard diet, yet PER was significantly lower in APOE4 flies housed on HSD compared to APOE3 flies on HSD (Fig 5E). Quantification of the response to 50mM fructose reveals a significant decrease in PER in APOE4 flies fed HSD compared to all other groups (Fig S4C). We further investigated the effects of HSD on aversive taste memory (Fig 5F). At 5 days of age there were no differences in responsiveness between APOE flies housed on standard diet or HSD (Fig 5G). Conversely, memory was significantly poorer in APOE4 flies fed a HSD compared to APOE3 flies receiving the same treatment at 20 days of age (Fig 5H). Twenty-day-old APOE4 flies fed with HSD showed a significant memory loss compared to all other groups (Fig S4D). Therefore, APOE4 flies are more sensitive to HSD across a number of behavioral assays, supporting the notion that APOE4 increases sensitivity to multiple different environmental stressors.

**Figure 5.**
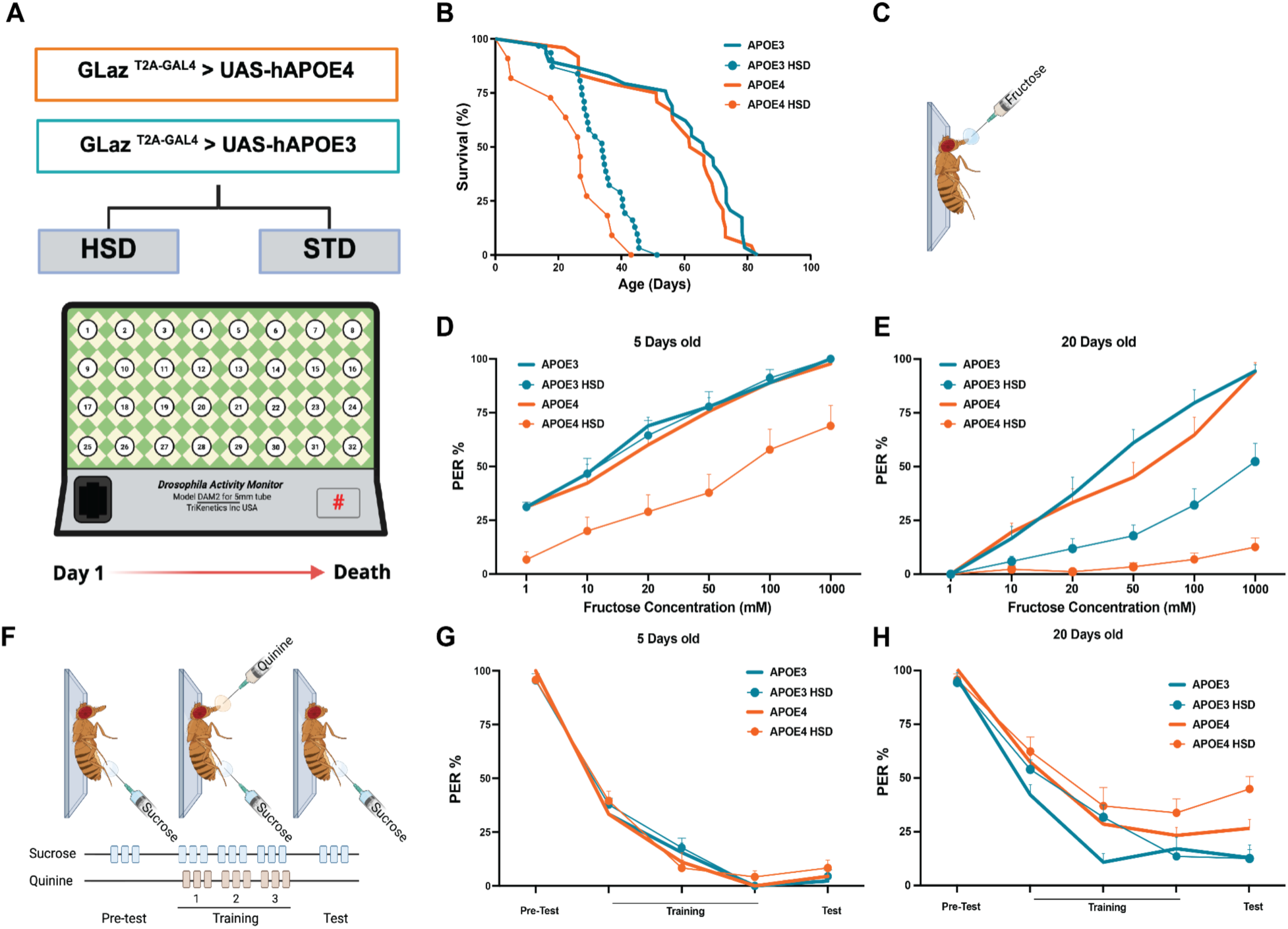
HSD reduces longevity and alters taste behavior in APOE3 and APOE4 flies. **(A)** Experimental design used to assess the effect of HSD on longevity. APOE3 or APOE4 flies were maintained on standard or HSD diet, and survival was monitored using the Drosophila Activity Monitor system. **(B)** HSD reduced survival in both APOE3 and APOE4 flies compared to their respective STD controls (log-rank Mantel-Cox test, χ²(3) = 75.21, *p* < 0.001; Gehan–Breslow-Wilcoxon test, χ²(3) = 56.19, *p* < 0.001). **(C)** Schematic of Proboscis Extension Reflex (PER) experiment. **(D)** At 5 days of age, HSD fed APOE4 flies exhibited significantly reduced PER compared with all the other groups (*p* < 0.001), whereas HSD did not significantly alter PER in APOE3 flies (*n* = 15)**. (E)** At 20 days of age, significant effects were detected for APOE genotype (*p* < 0.001) and diet (*p* < 0.001) and HSD reduces PER, with a significantly greater overall effect in APOE4 than APOE3 flies (*n* >16). **(F)** Schematic of aversive taste memory assay**. (G)** There were no significant differences between the groups at 5 days of age. **(H)** At 20 days of age, significant effects were detected for APOE, (p < 0.001), and diet (p = 0.0168) where genotype and diet alter the overall aversive-learning/memory trajectory, with APOE4-HSD flies showing the weakest memory at the final Test section (*n* > 16). Data are shown as mean ± SEM.

## Discussion

Here, we characterized multiple aging-associated phenotypes to define changes in health span in APOE model flies. The relatively short lifespan of *Drosophila* provides a powerful system for aging research, allowing age-dependent traits to be assessed across the lifespan^59^. Our findings indicate that several markers of aging, including declines in chemosensory function and memory, as well as increased gut permeability, are exacerbated in APOE4 flies exposed to ROS-inducing stress. These phenotypes mirror several age-related impairments that are commonly observed in humans carrying the AD-associated APOE4 allele ^4,7^. Together, these findings establish humanized APOE flies as a genetically tractable model for dissecting how APOE4 interacts with environmental and metabolic stressors to promote age-related functional decline and AD-relevant pathogenesis.

In this study we employed a humanized APOE model that complements widely used *Drosophila* models of Alzheimer’s disease. Most fly studies have relied on overexpression of pathogenic proteins such as Aβ42 or hyperphosphorylated Tau, which model downstream disease pathology ^46,60,61^. In contrast to these models that express causative protein variants, APOE4 is a common risk allele that increases disease susceptibility through interactions with aging and environmental factors. The fly model represents a T2A-GAL4 humanization strategy. Human APOE3 or APOE4 variants are expressed in place of the endogenous fly ortholog *GLaz* under native regulatory control, allowing direct comparison of APOE3 and APOE4 under physiologically relevant conditions ^27^. *GLaz* is broadly expressed in glial cells, as well as a subset of neurons, providing the opportunity to examine how APOE variants influence cellular function within the endogenous tissues that normally mediate lipid transport and homeostasis ^62^. Unlike transgenic overexpression models, this approach more closely recapitulates the subtle genetic effects associated with human APOE4, enabling the investigation of gene-by-environment interactions that are thought to underlie late-onset AD. Combined with the short lifespan and genetic tractability of *Drosophila*, this system provides a powerful platform for defining the cellular and behavioral consequences of APOE4 expression and for identifying interventions that enhance resilience to disease-associated stressors.

In this study we find that APOE4 flies are more sensitive to both rotenone and a high-sugar diet, supporting a model in which APOE4 increases vulnerability to physiological stress rather than acting as an isolated cause of disease ^63^. This is consistent with human studies showing that AD risk is strongly influenced by comorbid conditions, including obesity, type II diabetes, insulin resistance, cardiovascular disease, sleep disruption, and traumatic brain injury, many of which are associated with oxidative stress, metabolic dysfunction, inflammation, and impaired lipid homeostasis ^64,65^. The enhanced sensitivity of APOE4 flies to both ROS-inducing and metabolic stressors provides a tractable framework for testing how additional AD-associated challenges interact with APOE genotype and is consistent with a multi-hit model for AD associated with the APOE4 mutation in humans ^63^. Future studies could examine stressors such as chronic sleep loss, circadian disruption, inflammatory activation, traumatic brain injury, or exposure to additional environmental toxins. For example, prior work in Aβ42 mutant flies has shown that high-sugar diet disrupts phagocytic function, drawing parallels to microglial dysfunction observed in AD^66,67^. Therefore, humanized APOE flies provide a scalable system to test how genetic susceptibility combines with environmental and physiological stressors to drive age-dependent neurodegeneration.

A growing body of evidence suggests that impaired lipid handling and oxidative stress are central drivers of APOE4-associated neurodegeneration ^68,69^. APOE functions as a lipid transport protein that facilitates the movement of lipids between cells, while additional AD risk genes, including ABCA-family transporters and endocytic regulators such as PICALM and CD2AP, participate in pathways that promote the sequestration of oxidized lipids within glia^70^. Disruption of these pathways impairs glial lipid droplet formation, resulting in the accumulation of toxic lipid species, increased oxidative damage, and progressive neuronal dysfunction. In both *Drosophila* and mammalian systems, glial lipid droplets are thought to serve a neuroprotective role by buffering neurons from reactive oxygen species (ROS)-induced lipid toxicity ^71^. Consistent with this model, APOE4 flies in our study displayed enhanced vulnerability to ROS-inducing rotenone and to metabolic stress induced by a high-sugar diet. Rather than causing overt pathology under baseline conditions, APOE4 appears to reduce cellular resilience, rendering animals less capable of coping with oxidative and metabolic challenges that emerge during aging. The progressive deficits in memory, chemosensory function, gut integrity, and longevity observed in APOE4 flies are therefore consistent with a model in which impaired lipid homeostasis and ROS management drive age-dependent functional decline. These findings support the hypothesis that APOE4-mediated disruption of neuroprotective lipid-handling pathways contributes to disease susceptibility and suggest that humanized APOE flies provide a useful platform for dissecting the cellular mechanisms linking oxidative stress, lipid metabolism, and neurodegeneration.

Here, we broadly expressed APOE under control of the GLaz promoter. However, it is possible that APOE4, and other AD-associated variants, differentially affect specific neuronal and glial subtypes. For example, selective expression of Aβ42-Arctic in the mushroom bodies, a central brain region involved in learning, memory, and sleep regulation, induces neuronal hyperexcitability and disrupts sleep, suggesting that AD-associated variants can directly alter neural circuit function ^20^. Conversely, expression of Aβ42-Arctic in gustatory neurons results in loss of innervation to primary taste-processing centers and reduced neural responses to sugar stimulation, demonstrating that AD pathology can differentially impact distinct sensory circuits ^24^. Therefore, defining the cell-type-specific effects of APOE variants may provide important mechanistic insight into how APOE4 promotes disease susceptibility. Combining targeted expression approaches with functional imaging and longitudinal analyses across the lifespan could reveal how APOE variants alter neuronal and glial function during aging and identify the cellular populations most vulnerable to AD-associated pathology.

Multiple compounds have been identified in Drosophila models that ameliorate Aβ42-induced pathology. For example, treatment with the GABA-A receptor agonist gaboxadol restored memory deficits in flies expressing Aβ42-Arctic ^23^. Similarly, treatment with Levetiracetam reduced neuronal hyperexcitability and improved behavioral phenotypes in Drosophila models of neurodegeneration^20^. The identification of multiple APOE4-associated phenotypes in our model provides an opportunity to screen for compounds that selectively modify APOE4-dependent pathology. FDA-approved drug libraries and small-molecule screens have been successfully employed in Drosophila to identify modulators of sleep, feeding, and lifespan ^72–75^. Applying similar approaches to restore sensory function, cognitive performance, or longevity in APOE4 flies may provide a powerful avenue for identifying novel therapeutic strategies that protect against AD.

Taken together, these studies provide evidence that flies harboring a substitution of human APOE4 in place of the Drosophila ortholog *GLaz* provide a model for studying AD. This system recapitulates key aspects of APOE4-associated vulnerability and offers a tractable platform for identifying pharmacological interventions that restore function, defining genetic factors that modify APOE4-dependent phenotypes, and elucidating the cellular and molecular mechanisms that underlie disease susceptibility. By combining genetic, behavioral, and physiological approaches, this model has the potential to reveal conserved pathways linking APOE4 to neurodegeneration and to identify novel therapeutic strategies for AD

## Methods

### Fly Stocks and Husbandry

Flies were reared and maintained on a 12:12 light-dark cycle in humidified incubators at 25°C and 65% humidity (Percival Scientific, Perry, IA, USA). The fly stocks, GLaz ^T2A-GAL^^4^ (BDSC #77899), UAS-APOE3 (BDSC #76605); and UAS-APOE4 (BDSC #76607) were used in this study. F1s were generated by crossing female GLaz ^T2A-GAL^^4^ to UAS-APOE lines, creating flies expressing human APOE variants in a Glaz-specific expression pattern. All flies were reared on standard molasses fly food ^76^.

### Generation of rotenone and high-sugar diets

Rotenone was added to standard molasses-based food at a final concentration of 50 uM. A 20 mM stock solution was made by dissolving 7.8 g rotenone into 1 L DMSO and 2.5 uL of the 20 mM solution was added per 1 mL of fly food. To make high-sugar diet food, 30% extra sucrose was added to standard molasses-based food.

### Sleep Analysis

Sleep behavior was measured using the Drosophila Activity Monitoring System (DAM2; TriKinetics Inc., Waltham, MA), which detects locomotor activity based on infrared beam crossings of individual flies. Flies were briefly anesthetized with CO₂ and loaded into 65 mm × 5 mm glass locomotor tubes containing standard molasses based food or the specified dietary intervention (e.g., Rotenone/DMSO-supplemented food). Flies were acclimated for a minimum of 24 hours prior to the start of behavioral analysis. Food was placed at one end of each tube and was sealed with cotton plugs. Unless otherwise noted, all experiments were performed with 2-4 day-old mated male flies, which were selected to minimize age- and sex-dependent variability in sleep behavior. For lifespan-associated studies, sleep was assayed across different ages, with flies maintained on standard food and transferred to fresh tubes every 5 days to prevent food desiccation and microbial contamination. For experiments using the Drosophila Activity Monitoring (DAM) system (TriKinetics, Waltham, MA, USA), sleep and waking activity were quantified from infrared beam crossings of individual flies. Raw activity files were retrieved using DAMFileScan (TriKinetics), and custom Python scripts were used to bin activity at 5-min resolution. Sleep was defined as periods of immobility lasting ≥5 min, and sleep traits—including total sleep, mean bout length, bout number, and waking activity were extracted using the Drosophila Sleep Counting Macro ^77–79^.

### Longevity Assay

Lifespan was quantified using the Drosophila Activity Monitoring System (DAM2; TriKinetics Inc.), which continuously records infrared beam crossings of individual flies. Flies were collected within 24 h of eclosion and maintained in mixed-sex groups for 2 days to allow mating. Male flies were then separated under brief CO₂ anesthesia and individually loaded into 65 mm × 5 mm glass tubes containing standard Bloomington fly-food (Nutri-Fly, Genesee Scientific). Tubes were maintained in Percival incubators (DR-36VL, Percival Scientific) under controlled environmental conditions (25 °C, 55–65% humidity, 12:12 h light-dark cycle). Flies were transferred to new tubes with fresh food every 5 days, at which point survival was scored. Flies that escaped or were injured during transfers were excluded from the analysis. These exclusions accounted for less than 5% of the total flies tested. The time of death for each fly was operationally defined as the point of last recorded waking activity (final beam break) in the DAM system. Lifespan was calculated as the number of days survived post-eclosion until this endpoint. Similarly, group longevity was quantified using male flies housed in groups of 10 that were transferred in food vials containing standard or rotenone food. Flies were transferred into fresh vials every 2-3 days until death. They were maintained under controlled environmental conditions (25 °C, 55–65% humidity, 12:12 h light-dark cycle).

### Proboscis Extension Reflex (PER)

Flies were starved for 14 hours before PER testing, following previously described protocols ^37^. Flies were anesthetized using CO_2_ and mounted on microscope slides (#12-550-15, Fisher Scientific) with only the head and proboscis exposed. After a 60-minute acclimation period in a humidified chamber, flies were offered water until satiated. A 1-ml ultrafine insulin syringe was used for tastant application. Solution was applied to the tip of the proboscis for 1–2 seconds. Full proboscis extensions were scored as positive responses. Each tastant was presented three times at 1-minute intervals, and PER was calculated as the percentage of positive responses per fly.

### Aversive taste memory assay

Taste discrimination was assessed by measuring aversive taste memory, as described previously ^51,80^. Mated male flies at 5 and 20 days of age were starved for 14 hr prior to each experiment. Flies were later anesthetized on CO_2_ pads, glued with clear nail polish (#451D; Wet n Wild, Los Angeles, CA) to dorsal side of thorax on a microscopy slide (StatLab^TM^ ColorView Microscope Slide) and were acclimated to these conditions in a humidified box for 60 min. For each experiment, the microscope slide was mounted vertically under a dissecting microscope (#SM-1BSZ-144S; AmScope, Irvine, CA). Flies were water satiated prior to each experiment and in between each test/training session, lasting ∼2 min. A 1-ml ultra fine insulin syringe was used for tastant presentation. We used purified water, 500 mM sucrose (Thermo Fisher Scientific) or 50 mM quinine (Sigma-Aldrich) solutions. For the pretest, each fly was given 500 mM sucrose on their tarsi three times with a 30 s interval between applications, and the number of full proboscis extensions was recorded. During training, a similar protocol was used except that each sucrose presentation was immediately followed by quinine presentation, in which flies were allowed to drink for up to 2 s or until an extended proboscis was retracted. A total of three training sessions were performed. To assess taste discrimination, flies were tested with sucrose only. At the end of each experiment, flies were given 1 M sucrose to check for retained ability to extend proboscis, and all non-responders were excluded.

### Blue-dye feeding assay

Short-term food consumption was measured as previously described ^55^. Flies were either fed or starved for 24 h on moistened Kimwipes or maintained on standard food. At ZT0, flies were transferred to vials containing 1% agar, 5% sucrose, and 2.5% FD&C Blue Dye No. 1. After 30 minutes of feeding, flies were flash-frozen on dry ice and homogenized individually in 400 μL PBS (pH 7.4). Absorbance at 655 nm was measured using a 96-well plate reader (Millipore iMark, Billerica, MA). Background absorbance was corrected by subtracting the mean absorbance of non-dye-fed control flies.

### Gut permeability Assay

Intestinal integrity was assessed using the Smurf assay, as previously described ^53^. First, freshly emerged flies were isolated and provided time to mate for 2 days. Male flies were then separated by anesthetizing with mild CO2 and placed into vials containing standard food at a density of ∼20 flies per vial. At ZT 0, flies of a given age and genotype were transferred onto fresh medium containing blue dye (2.5% w/v; FD&C blue dye #1) for 24 hrs. At ZT 0 the following day, the percentage of Smurf flies in each vial was recorded. Flies were considered Smurf if blue coloration extended beyond the gut.

### Olfactory Trap Assay

Groups of 30 male flies were collected and food-deprived for 6 h. Following starvation, flies were transferred to agar-containing bottles equipped with two microcentrifuge-tube traps. Each trap contained a small cotton plug saturated with either experimental (acetic acid, thermo fisher) or control solution (DI water) and was inserted into the agar with the opening accessible to the flies. Bottles were covered with mesh and maintained in an undisturbed location for 4 h. Flies were then anesthetized with CO₂, and the numbers of flies located inside each trap and remaining outside the traps were recorded.

### Statistical Analysis

Statistical analyses were performed using either RStudio (Version 2026.04.0+526) or GraphPad Prism (Version 10.5.0). For sleep and activity profile the data was analyzed using two-way ANOVA to test the effects of age and genotype/treatment group, followed by Tukey’s multiple-comparison test where needed. Longitudinal sleep measurements were analyzed across age with each group included only through its 50% survival point to reduce the noise from progressive mortality.

Longevity datasets were analyzed using Kaplan-Meier survival curves and differences between the genotypes and treatment conditions were analyzed using the log-rank (Mantel-Cox) test with multiple comparison testing where needed. Survival curves were also evaluated using the Gehan-Breslow-Wilcoxon test, and median survival with 95% confidence intervals were calculated for each group. Smurf assay data was analyzed using two-way ANOVA to assess the effects of genotype and treatment at the given age, followed by Tukey’s multiple comparison test.

For memory and PER assay, each fly was tested three times at each trial, and responses were shown as the proportion of positive trials. Data are shown as estimated response probabilities with 95% confidence intervals. Data were analyzed using a binomial generalized estimating equation with fly identity as the repeated-measures cluster and APOE genotype, food treatment, fructose concentration, and their interactions as fixed effects. Šídák’s multiple comparison test was used to compare treatment effects within each genotype and genotype effects within each treatment. Adjusted *p* < 0.05 was considered statistically significant.

For experiments involving two categorical factors, data were analyzed using two-way factorial models to test the main effects of each factor and their interaction. Experiments with age as an additional factor were analyzed using three-way factorial models. Significant effects were followed by multiple-comparison-adjusted pairwise tests as indicated in the corresponding figure legends.

## Acknowledgements

This work was supported by a gift from the WoodNext Foundation to MM and ACK, and NIH R01NS131628 to ACK.

## Supplemental Figures

**Figure S1.**
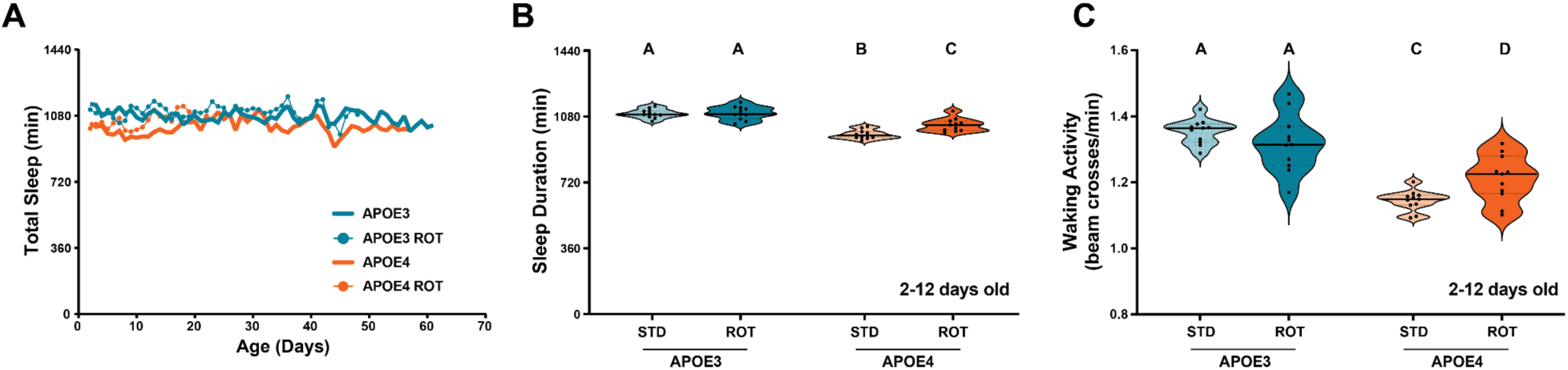
Sleep across lifespan. **(A)** Sleep profiles were analyzed only up to the 50% survival point for each group. Sleep duration was not significantly different across the lifespan among the groups**. (B)** Total sleep during days 2-12 was significantly lower in APOE4 flies, compared to APOE3 flies (two-way ANOVA, group effect: F_(3,1568)_ = 70.37, *p* < 0.001). Sleep duration was not affected by rotenone in APOE3 flies while significantly increased sleep duration in APOE4 flies (*p* < 0.01). **(C)** There was no effect of rotenone on waking activity in 2 to 12 days old APOE3 flies (*p* > 0.05). Conversely, waking activity was increased in rotenone-fed APOE4 flies (*p* < 0.001), (two-way ANOVA, group effect: F_(3,1578)_ = 39.48, *p* < 0.001).

**Figure S2.**
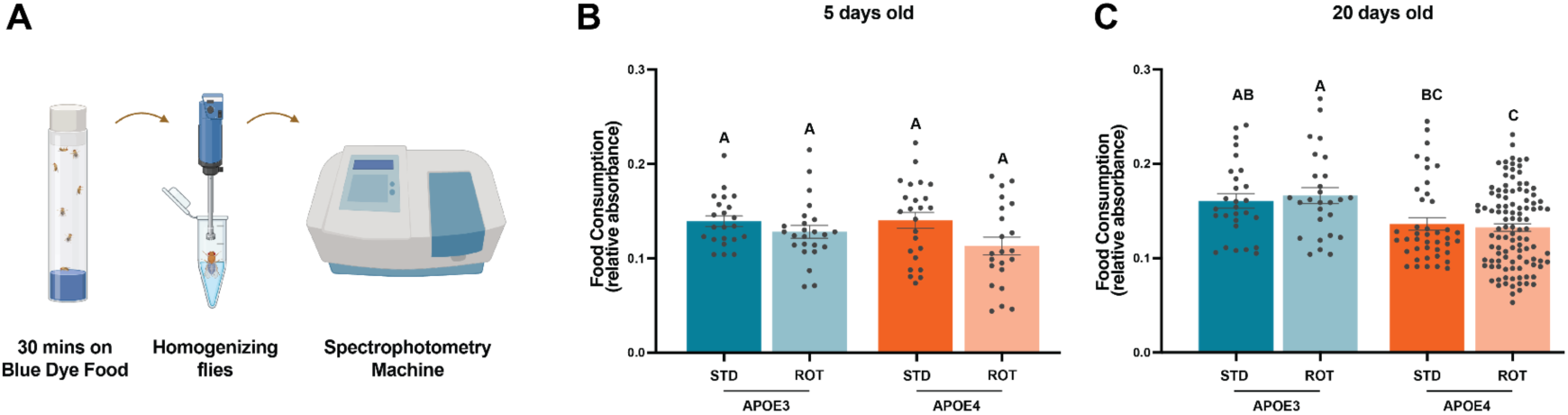
Comparison of feeding in APOE3 and APOE4 flies. **(A)** Starved flies were transferred to blue dye food for 30 minutes. After homogenizing them, spectrophotometry was used to quantify the amount of consumed food. **(B)** No significant differences in food consumption were detected among the flies at age of five (*n*>22). **(C)** In 20-day-old flies food consumption differed significantly among groups (one-way ANOVA, F_(3,200)_ = 6.774, *p* = 0.0002), though no significant differences were detected between DMSO and rotenone treated flies within either APOE3 or APOE4 genotypes (*n* = 28). All data are shown as mean ± SEM.

**Figure S3.**
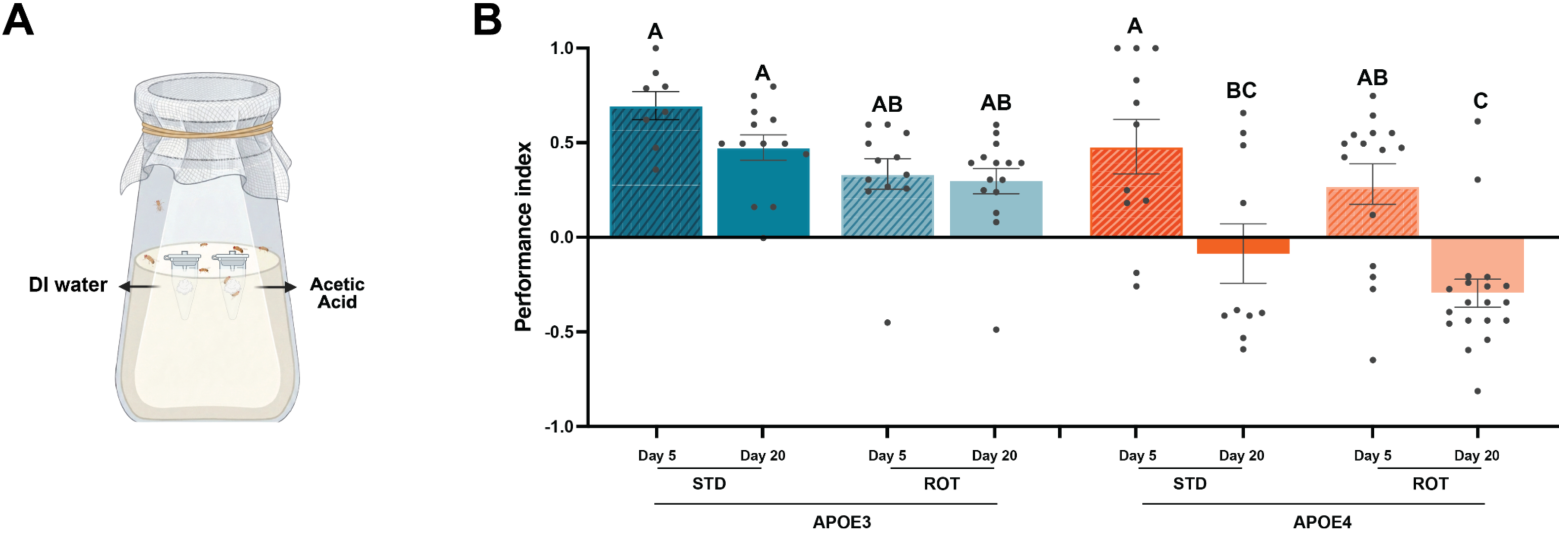
Olfactory trap assay in APOE3 and APOE4 flies. **(A)** Schematic of Trap assay. **(B)** Trap assay preference was measured in 5 and 20-day-old flies. No significant decrease was observed in olfactory performance of APOE3 flies, while there was no preferential selection of acetic acid in 20-day-old in APOE4 flies on rotenone or standard food (*p* < 0.05). Groups not sharing a letter differ at adjusted *p* < 0.05. 5-day (*n* > 8). All data are shown as mean ± SEM.

**Figure S4.**
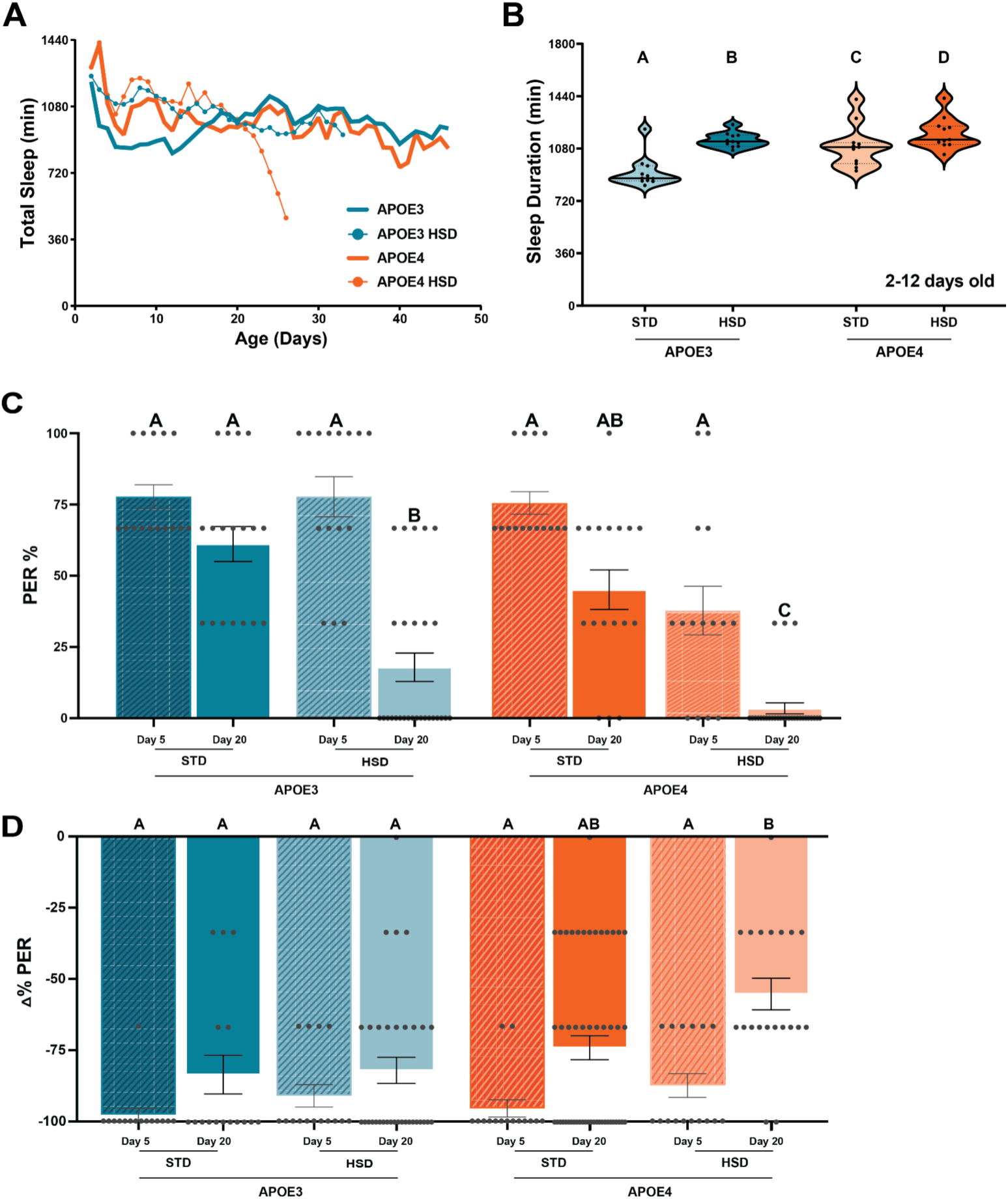
High sugar alters sleep and taste related behavioral phenotypes in APOE3 and APOE4 flies. **(A)** Longitudinal total sleep in APOE3 and APOE4 flies maintained on STD or HSD, with each group analyzed only up to its own 50% survival point. HSD increased total sleep significantly in both APOE3 and APOE4 flies compared to their standard fed control flies (both *p* < 0.001). **(B)** Total sleep during days 2-12 differed significantly among groups (two-way ANOVA, group effect: F_(3,896)_ = 197.2, *p* < 0.001). Tukey’s multiple-comparisons test showed increased sleep in both APOE3 and APOE4 flies on HSD compared with their standard food fed respective controls (both *p* < 0.001). **(C)** PER responses to 50 mM fructose were analyzed separately. 20-day old APOE4 flies on HSD displayed the lowest PER and differed significantly from each of the other groups (*p* < 0.05). **(D)** At the age of 20, HSD fed APOE4 flies showed the most negative ΔPER compared to other groups labeled A, whereas 20-day old standard food fed APOE4 flies showed an intermediate response that did not significantly differ from all of the other groups. Groups not sharing a letter differ at adjusted *p* < 0.05. Data are shown as mean ± SEM.

